# Silencing neuronal activity is required for developmental circuit remodeling

**DOI:** 10.1101/2021.10.31.466652

**Authors:** Oded Mayseless, El Yazid Rachad, Gal Shapira, André Fiala, Oren Schuldiner

## Abstract

Postnatal refinement of neuronal connectivity shapes the mature nervous system. Pruning of exuberant connections involves both cell autonomous and non-cell autonomous mechanisms, such as neuronal activity. While the role of neuronal activity in the plasticity of excitatory synapses has been extensively studied, the involvement of inhibition is less clear. Furthermore, the role of activity during stereotypic developmental remodeling, where competition is not as apparent, is not well understood.

Here we use the *Drosophila* mushroom body as a model to show that regulated silencing of neuronal activity is required for developmental axon pruning of the γ-Kenyon cells. We demonstrate that silencing neuronal activity is mechanistically achieved by cell autonomous expression of the inward rectifying potassium channel (irk1) combined with inhibition by the GABAergic APL neuron. These results support the Hebbian-like rule ‘use it or lose it’, where inhibition can destabilize connectivity and promote pruning while excitability stabilizes existing connections.

## Introduction

The precise connectivity between neurons determines how the brain perceives and processes information. The mature connectivity of the nervous system is often established by refinement of exuberant connections that were established during embryonic development (reviewed in: Akin and Zipursky, 2020; Luo and O’Leary, 2005; Schuldiner and Yaron, 2014). Such refinement of neuronal connectivity, also known as neuronal remodeling, is a widespread and conserved strategy to sculpt neuronal circuits. Neuronal remodeling is governed by a combination of cell-autonomous genetic programs (Luo and O’Leary, 2005; Riccomagno and Kolodkin, 2015; Yaniv and Schuldiner, 2016), as well as non-cell-autonomous processes (Hensch, 2005; Katz and Shatz, 1996; Meltzer and Schuldiner, 2020). One of the major extrinsic factors that influence refinement and remodeling of connectivity is neuronal activity. Indeed, experience-dependent competition has been shown to sculpt neuronal circuits undergoing remodeling in various systems, spanning the neuromuscular junction to establishment of connectivity in sensory systems (Chung and Barres, 2009; Hashimoto et al., 2009; Johnson-Venkatesh et al., 2015; Katz and Shatz, 1996; Tapia et al., 2012; Walsh and Lichtman, 2003). However, how inhibitory activity influences refinement of neuronal connectivity or how neuronal activity influences remodeling in systems in which neuronal competition is not clearly apparent, is not fully understood.

The mushroom body (MB) is a neuronal structure in insects that is implicated in associative learning and memory (Bilz et al., 2020; Heisenberg et al., 1985), and receives mostly olfactory input from the antennal lobe (Takemura et al., 2017; Zheng et al., 2018). The MB circuit includes intrinsic MB neurons, called Kenyon cells (KCs), several modulatory neurons, such as the GABAergic anterior paired lateral neuron (APL) and distinct dopaminergic neurons (DANs), as well as the MB output neurons (MBONs). The MB forms a functional neural circuit during larval life, which then remodels to give rise to the adult MB circuit (Lee et al., 1999; Technau and Heisenberg, 1982). How the entire circuit remodels during metamorphosis is unclear. For many years it has been appreciated that the first-born KCs, the γ-KCs, undergo stereotypic remodeling that involves pruning of larval axons and dendrites followed by regrowth to form adult connections (Watts et al., 2003). Recently, we found that the APL neuron also undergoes massive rearrangement of its axons and dendrites during metamorphosis (Mayseless et al., 2018). Interestingly, the remodeling of the γ-KCs and the APL is coordinated, at least in part via neuronal activity in the γ-KCs and calcium/calmodulin signaling within the APL (Mayseless et al., 2018). However, whether and how neuronal activity, i.e. membrane depolarization and subsequent transmitter release, directly affects remodeling of the different circuit components is not known.

Here, we test a hypothesis that expands the Hebbian based plasticity hypothesis by suggesting that reduced neuronal activity is required for axon pruning. Indeed, we found that silencing activity, likely by lowering membrane potential, is a prerequisite for the initiation of pruning. We suggest that γ-KCs achieve this by two non-mutually exclusive mechanisms: the cell autonomous upregulation of the inward rectifying potassium channel Irk1 expression, and inhibitory inputs relayed by the GABAergic APL neuron.

## Results

### Calcium levels in γ-KCs are dynamic upon transition into metamorphosis

Studies from several neural systems have demonstrated that neuronal activity and Ca^2+^ signaling play a vital role in the coordination and control of neuronal refinement (Golovin et al., 2019; Kanamori et al., 2013; Kano et al., 2018; Mayseless et al., 2018; Qiu et al., 2020). We therefore set out to examine the Ca^2+^ dynamics during the stereotypic remodeling of γ-Kenyon cells (KCs; Figure 1 A, Lee et al., 2000), the major intrinsic cell type within the *Drosophila* mushroom body (MB). While monitoring Ca^2+^ dynamics using genetically encoded calcium indicators (GECI) such as GCaMP offers excellent sensitivity and resolution, it is less optimized for exploring long term changes in Ca^2+^ levels in deep tissues such as the MB. CaMPARI is an engineered ratiometric fluorescent protein, which undergoes efficient and irreversible green-to-red conversion only when elevated Ca^2+^ and experimenter-controlled illumination coincide (Fosque et al., 2015; Moeyaert et al., 2018). Thus, CaMPARI offers the possibility to image Ca^2+^ dynamics over a relatively long period, and to compare relative activity levels between different developmental stages. Using this tool, we examined relative Ca^2+^ levels in γ-KCs before and at the onset of remodeling - between early 3^rd^ instar larvae (L3), and pupae up to 6 hours after puparium formation (h APF). We observed a significant decline in relative Ca^2+^ levels at 0h APF, the onset of metamorphosis, as compared to larval stages (Figure 1 B – 0h APF one-Sample Wilcoxon Signed Rank Test, with Bonferroni correction, p=0.0024). Subsequently, and to our surprise, Ca^2+^ levels started increasing at 3h APF and reached elevated levels compared to larval stages (Figure 1 B – 6h APF one-Sample Wilcoxon Signed Rank Test, with Bonferroni correction, p= 4.88281e^-4^). These results demonstrate that γ-KC calcium levels are highly dynamic during the transition from larva to pupa, even in the presumed absence of external inputs.

**Figure 1.**
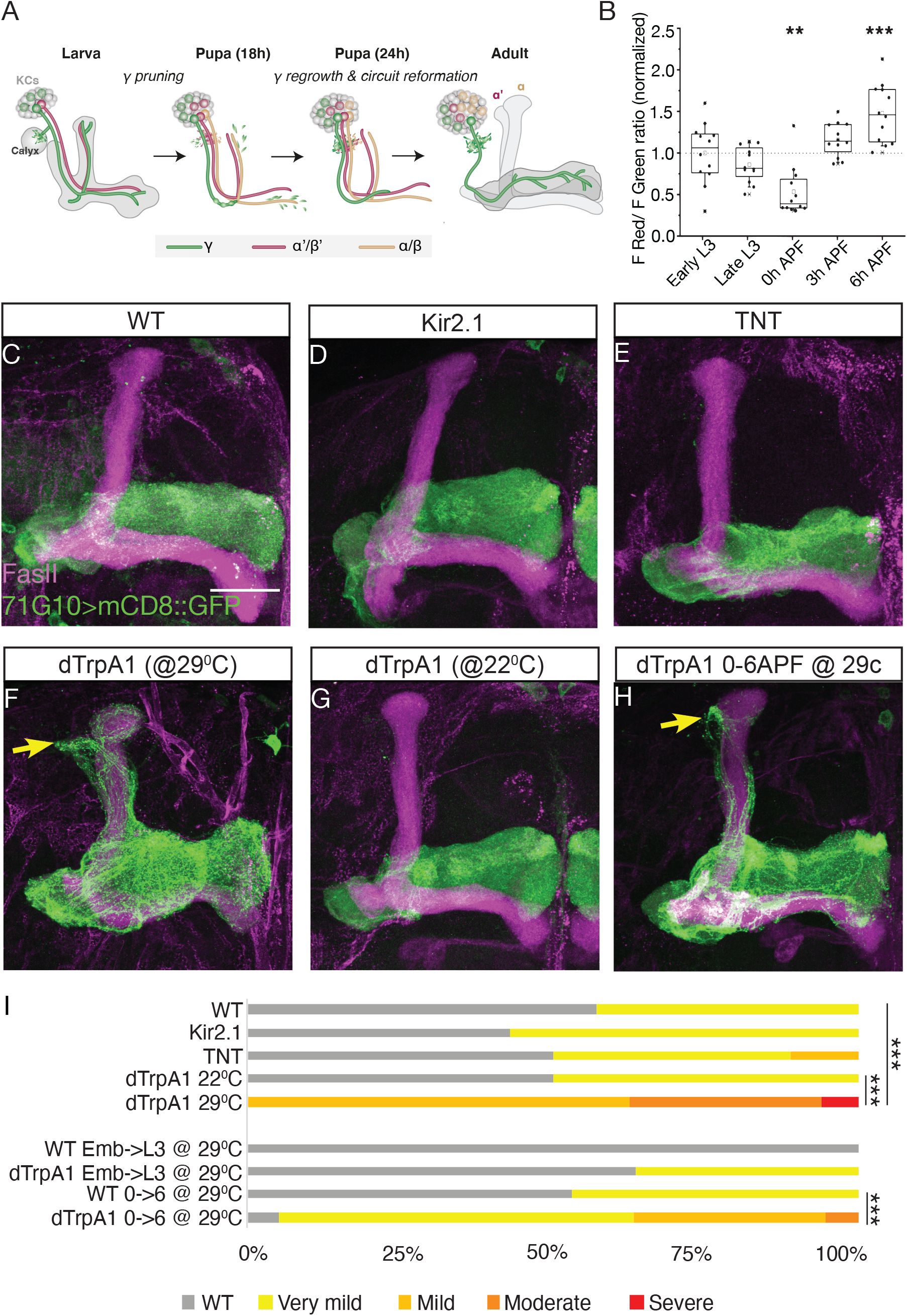
Chronic activation of γ-KCs during early pupal stages inhibits pruning. **(A)** Cartoon illustration depicting developmental remodeling of γ-KCs. Larval γ-KCs send dendrites to the MB calyx and a bifurcated axon which forms the MB lobes. During metamorphosis γ-KC dendrites and axons prune up to a set point, and later regrow to form adult specific connections. **(B)** Normalized red to green fluorescence ratio of CaMPARI2.L398T driven under the control of the R71G10-Gal4 driver at the indicated times. L3 is 3^rd^ instar larvae, APF is after puparium formation. Late vs early L3 larval stage was set by lack of gut coloration. Statistical significance calculated using one-Sample Wilcoxon Signed Rank Test, with a Bonferroni correction and is, p=0.0024 for 0h APF and p<0.0001 for 6h APF compared to early L3. Box plots indicate median values (lines), mean values (open squares), inter-quartile ranges (boxes), 10/90 percentiles (whiskers) and individual data points (black squares). All data points are normalized to the mean of early L3 larvae **(C-H)** Confocal Z projections of adult brains immunostained with anti-FasII (magenta) and expressing (C) mCD8::GFP (green) under the control of GMR71G10-Gal4, or additionally expressing (D) Kir2.1, (E) TNT, or (F-H) dTRPA1 and grown at different temperatures (F) 29^0^C (G) 22^0^C (H) 29^0^C from 0-6h APF. **(I)** Ranking score quantification of pruning defect severity, as exemplified in Figure 1S. Statistical significance calculated using a Mann Whitney U test and is p<0.001 for dTrpA1 at 29^0^C (F, n=16) vs trpA1 at 22^0^C (G, n=6); p<0.001 for dTrpA1 at 29^0^C (F, n=16) vs WT (C, n=14); and p<0.001 for pupa raised at 29^0^C from 0-6h APF with dtrpA1 (H, n=19) vs the same treatment for WT lacking dTrpA1 (not shown, n=17). Scale bar indicates 30*µ*m.

### Chronic activation of γ-KCs during key stages of remodeling inhibits pruning

Intracellular calcium transients often reflect changes in membrane potential, suggesting that γ-KCs neuronal excitability may play a role in the progression of neuronal remodeling. To determine whether depolarization or hyperpolarization of γ-KCs is required for pruning, we manipulated their activity via genetically encoded transgenes (Venken et al., 2011b) and examined the effect on their pruning. Hyperpolarizing γ-KCs, by expressing the inward rectifying K^+^ channel Kir2.1 (Baines et al., 2001) did not affect pruning (Figure 1 C-D, quantified in I, see S1A-E for ranking examples). Additionally, inhibiting neurotransmission, by expressing tetanus toxin light chain (TNT) in γ-KCs, did not affect pruning (Figure 1E, quantified in I). In contrast, activating γ-KCs by expressing the thermo-sensing cation dTrpA1 channel (Rosenzweig et al., 2005), resulted in a dramatic inhibition of pruning at the permissive 29°C as compared to the restrictive 22°C (Hamada et al., 2008) (Figure 1 F compared to G quantified in I, Mann Whitney U test, p=0.0015 vs trpA1 at 22^0^C and 1.6e^-05^ vs WT). These data suggest that chronic activation of γ-KCs inhibits their pruning.

Next, we explored the temporal window in which γ-KC activation affected pruning. We took advantage of the temperature sensitivity of the dTrpA1 channel to induce neuronal activation during different stages of development. Interestingly, raising flies in 29°C (resulting in the opening of the dTrpA1 channel) for the duration of their larval life, and transferring them to 22^0^C at 0h APF did not inhibit pruning of γ-KCs. In contrast, transferring flies to 29°C from 0 to 6h APF significantly inhibited pruning (Figure 1 H, quantified in I, Mann Whitney U test, p=0.0040).

To verify that chronic opening of dTrpA1 indeed induces chronic activation of γ-KCs, we used CaMPARI to examine the Ca^2+^ levels of pupal γ-KCs expressing dTrpA1. As expected, Ca^2+^ levels were significantly elevated upon dTrpA1 expression (Figure 1S F-G). Together, these results suggest that activation of γ-KCs and elevated Ca^2+^ levels at the onset of metamorphosis are sufficient to inhibit pruning, implying that hyperpolarization of γ-KCs at 0h APF might be required for their pruning.

**Figure S1.**
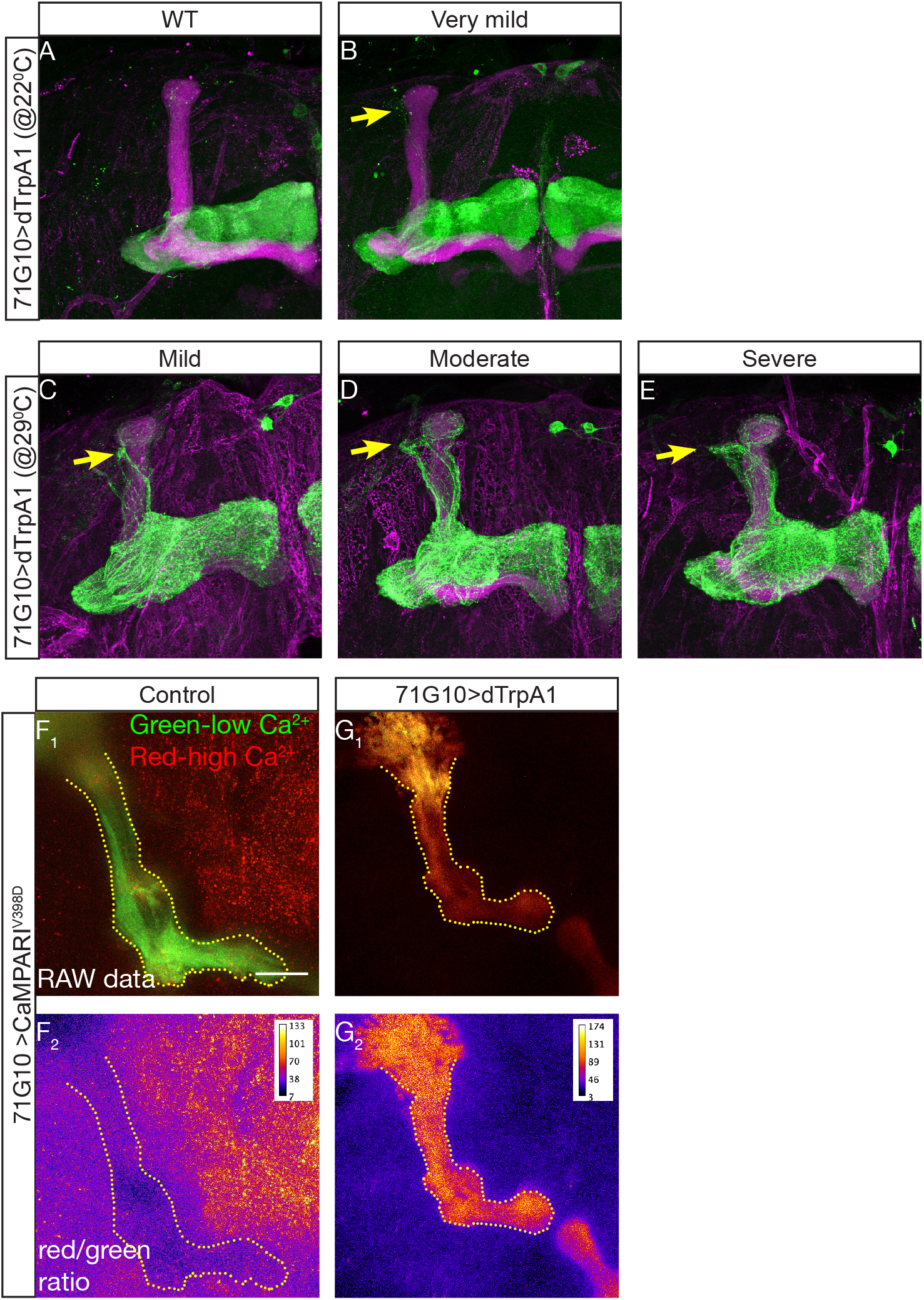
Examples of ranked pruning quantifications and dTrpA1 CaMPARI validation. **(A-E)** Examples of the ranking of pruning defect severity in experiments shown in Figure 1. Yellow arrowheads point towards unpruned axons. **(F-G)** Confocal Z projections of adult brains reared at 29^0^C, expressing CaMPARI^V398D^ (F), or dTRPA1 and CaMPARI^V398D^ (G), under the control of GMR71G10-Gal4 and exposed to photoconverting light for 15 minutes at 6h APF. (F^1^ and G^1^) Green indicates low relative Ca^2+^, while red indicates high relative Ca^2+^. (F_2_ and G_2_) ratio of red to green fluorescence. Dotted yellow line outlines MB structure. Scale bar indicates 30*µ*m.

### Hyperpolarization of γ-KCs is required for pruning

Opening of dTrpA1 channels can have a dual effect; ion influx into γ-KCs which would induce membranal depolarization, and a subsequent release of neurotransmitters from the presynaptic sites. In order to examine which of these mediated the inhibition of pruning, we co-expressed dTrpA1 either with mammalian Kir2.1 to inhibit membrane depolarization, or with Tetanus toxin (TNT) to prevent exocytosis of evoked neurotransmitter containing vesicles. While co-expression of either Kir2.1 or TNT suppressed the dTrpA1-induced pruning defect to a significant degree, Kir2.1 suppression was significantly more penetrant (Figure 2 A-C, quantified in D, using Mann Whitney U test, p-value =0.03752 for dTrpA1,mCD8, n=13 vs dTrpA1,TNT n=22, and p<0.00001 for dTrpA1,mCD8 vs dTrpA1,Kir2.1, n=28, and p-value=0.00064 for dTrpA1,kir2.1 vs dTrpA1,TNT). These results suggest that the primary effect of dTrpA1 opening, in the context of γ-KC pruning, is the depolarization of γ-KC membranes rather than the secretion of neurotransmitters; yet it does not completely rule out non-cell autonomous influence. Taken together, these results indicate that hyperpolarization of γ-KCs is required for the initiation of their pruning.

**Figure 2.**
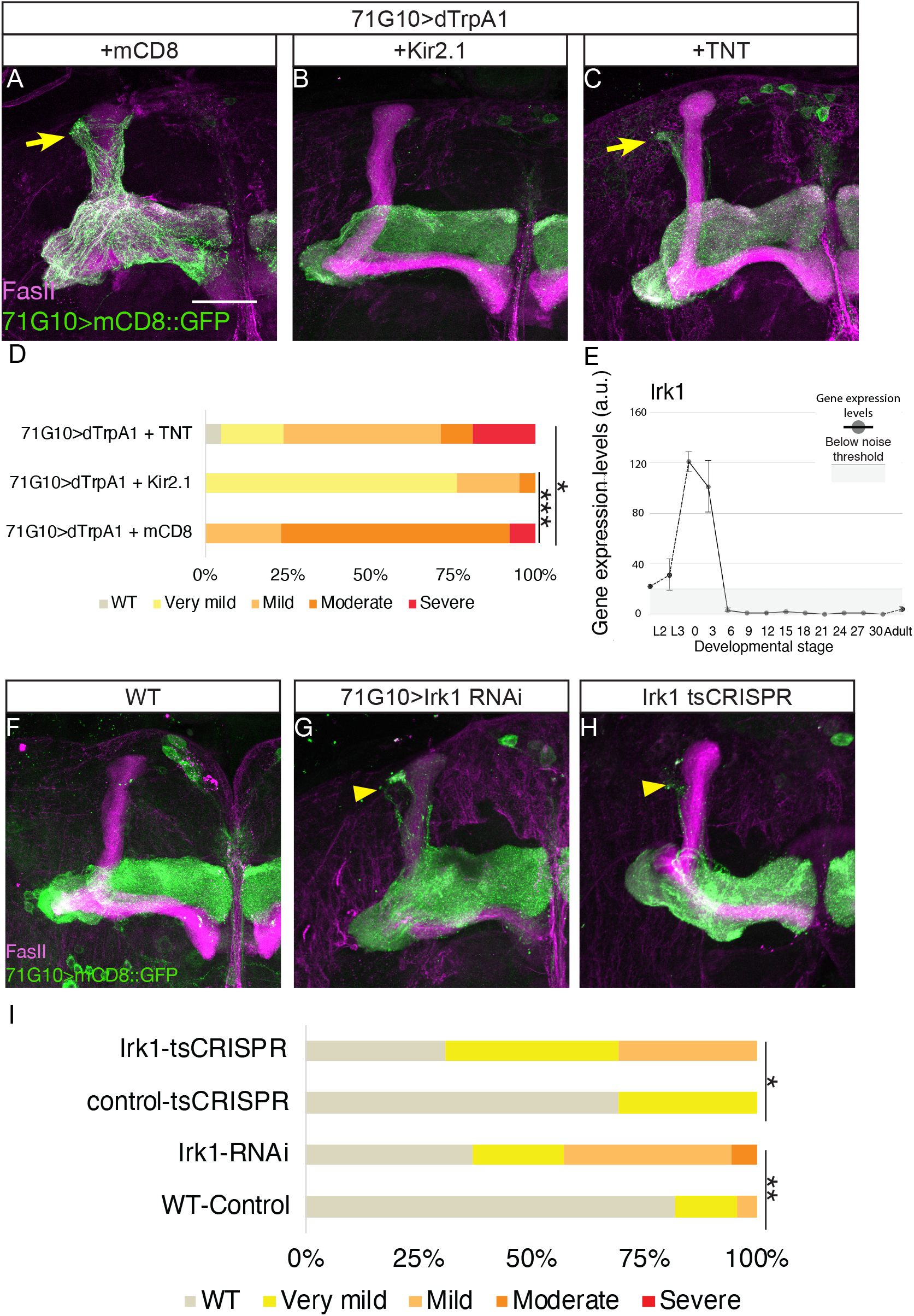
Hyperpolarization of γ-KC is required for pruning. **(A-C)** Confocal Z projections of adult brains reared at 29^0^C, immunostained with anti-FasII (magenta) expressing mCD8::GFP and dTRPA1 under the control of GMR71G10-Gal4, additionally expressing (A) an additional copy of mCD8::GFP, (B) Kir2.1, or (C) TNT. **(D)** Ranking of pruning defects in A-C. Statistical significance calculated using Mann-Whitney test, and is p-value =0.03752 for dTrpA1, TNT (C, n=22) vs dTrpA1, mCD8, (A; n=13) and p< 0.001 for dTrpA1, Kir2.1, (B; n=28) vs the control (A). **(E)** Expression profile of Irk1 in γ-KCs throughout development, demonstrating an acute increase in expression levels at the onset of metamorphosis. Extracted from (Alyagor et al., 2018), L2-L3 are 2^nd^ and 3^rd^ larval stages, 0-30 are hours APF. **(F-H)** Confocal Z projections of brains immunostained with anti FasII (magenta), expressing mCD8::GFP under the control of GMR71G10-Gal4 (F), or additionally expressing UAS-Irk1-RNAi^HMS02480^ (G), or UAS-Cas9.C and U6.2:Irk1-gRNA (H). **(I)** Ranking of pruning defects in F-H. Statistical significance calculated using Mann-Whitney test, p-value = 0.00168 for Irk1 RNAi (G; n=22) vs control (F; n=35) and 0.01046 for tsCRISPR (H; n=13) vs control (not shown; n=39). Scale bar indicates 30*µ*m.

To identify potential cell-autonomous mechanisms through which γ-KCs could hyperpolarize, we examined their developmental transcriptional landscape (Alyagor et al., 2018). Interestingly, we identified *inwardly rectifying potassium channel 1* (*irk1*) as specifically upregulated in γ-KCs at the onset of metamorphosis (Figure 2 E). In mammals, Irk channels participate in important cellular functions such as control of the resting membrane potential, maintenance of K^+^ homeostasis, and transduction of cellular metabolism into excitability (Döring et al., 2002). Indeed, perturbing the expression of Irk1 using RNAi or tissue-specific (ts)CRISPR (Meltzer et al., 2019) inhibited γ-KCs pruning (Figure 2 F-H quantified in I, using Mann Whitney U test, p-value = 0.00168 for Irk1 RNAi n=22 vs control n=35 and 0.01046 for tsCRISPR n=13 vs control n=39).

Overall, these results suggest that hyperpolarization of γ-KCs is required for their pruning, and demonstrate that cell-autonomous expression of Irk1 at the onset of metamorphosis is required for pruning, potentially by contributing to γ-KC hyperpolarization.

### APL activity and GABA-B-R1 expression are required for efficient γ-KC pruning

While Irk1 expression is required for γ-KC pruning, the pruning defect caused by its perturbation was much milder than that induced by chronic activation of γ-KCs (via TrpA1). Therefore, we set out to examine whether inhibitory neural signaling was also involved. The sole inhibitory input to γ-KCs before metamorphosis is considered to be the GABAergic anterior paired lateral (APL) neuron (Masuda-Nakagawa et al., 2014). Moreover, we have recently shown that APL remodeling is coordinated with that of γ-KCs (Mayseless et al., 2018). Therefore, we hypothesized that APL neuronal activity is a prime suspect to relay inhibitory signals to γ-KCs at these stages.

To test the role of APL activity in γ-KC pruning, we silenced the APL by expressing Kir2.1 and examined the concurrent effects on γ-KCs. Due to the stochastic nature of the APLi driver (Lin et al., 2014), we could analyze brains in which the APL is labeled and manipulated only in one brain hemisphere, while the second hemisphere remains unperturbed. Indeed, hemispheres in which the APL neuron expressed Kir2.1 displayed a mild yet significant γ-KC pruning defect compared to control hemispheres (Figure 3 A-C quantified in J, using paired Wilcoxon signed-rank test with continuity correction, p=0.000133 for APL>Kir2.1, n=31). In addition, silencing APL neurotransmitter secretion by expressing tetanus toxin within the APL also inhibited γ-KC pruning (Figure 3 G-I quantified in J, using paired Wilcoxon signed-rank test with continuity correction, p= 0.02473, n= 14). Interestingly, expressing tetanus toxin in the APL throughout development also induced blebbing of the APL neurites in some of the brains (Figure S2-A-C red arrowheads) but whether or how this is related to γ-KC pruning remains to be investigated. These results therefore suggest that APL activity is required for effective γ-KC pruning. Consistent with our hypothesis that the APL confers a hyperpolarizing effect that promotes pruning in γ-KCs, silencing APL neuronal activity should result in increased excitability of the γ-KCs. To explore this potential epistasis, and to test whether the increased excitability of γ-KCs is the cause for their defective pruning, we simultaneously expressed Kir2.1 in the APL and also in the γ-KCs. Indeed, this suppressed the APL-Kir2.1 driven pruning defect (Figure 3 G-I-quantified in J, using paired Wilcoxon signed-rank test with continuity correction, p= 0.833, n=12).

**Figure 3.**
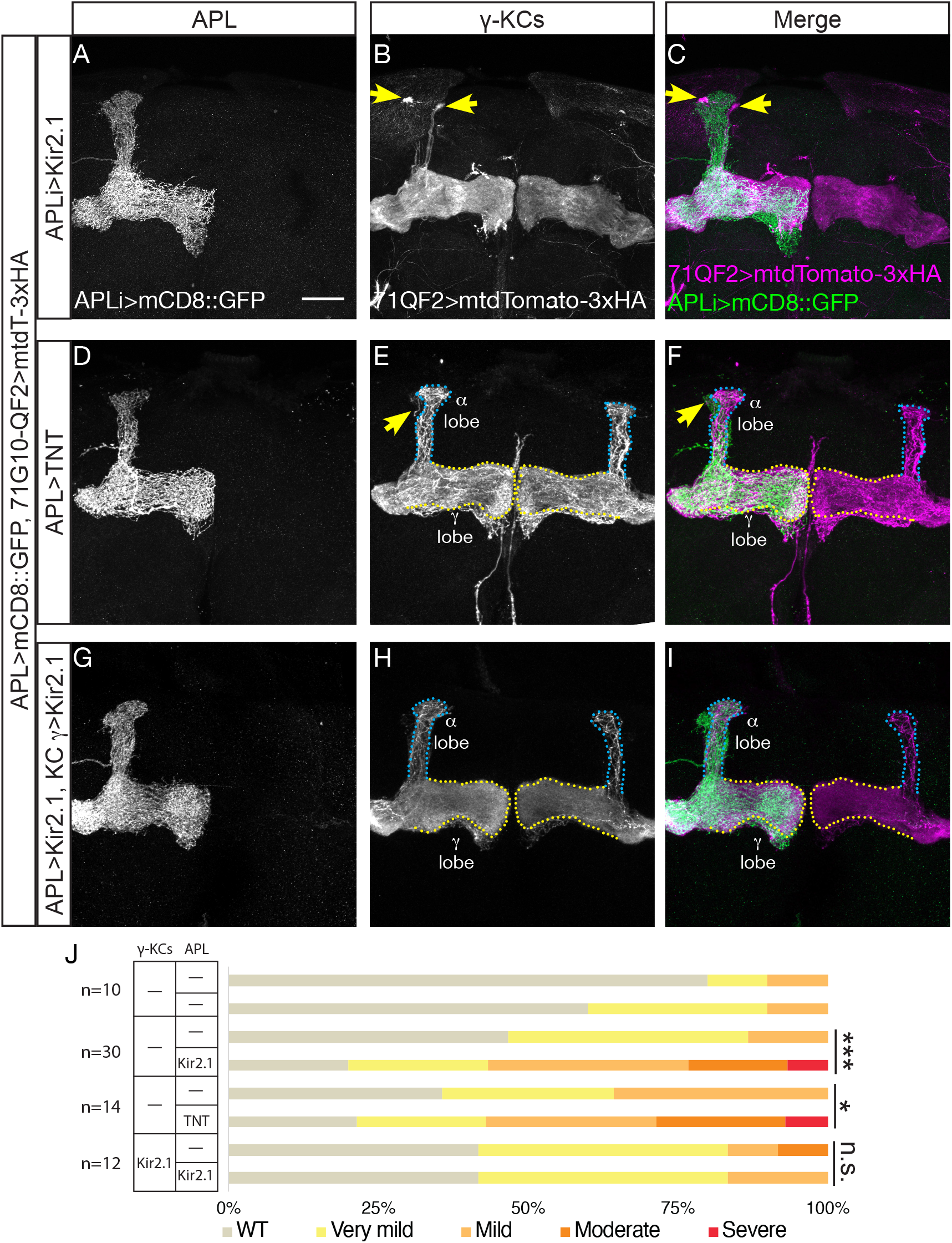
APL activity is required but not sufficient for γ-KCs pruning. **(A-I)** Confocal Z projections of brains expressing mCD8::GFP under the control APLi-Gal4 (grey in A,D,G and green in C,F,I) and mtdTomato-3xHA under the control of GMR71G10-QF2 (grey in B,E,H and magenta in C,F,I), additionally expressing Kir2.1 under the control of APLi-Gal4 (A-C), TNT under the control of APLi-Gal4 (D-F), or UAS-Kir2.1 under APLi-Gal4, as well as QUAS-Kir2.1 under GMR71G10-QF2 (G-I). Yellow arrowheads mark unpruned axons. Blue dotted outlines mark leaky expression of GMR71G10-QF2 in α/β KCs, and yellow dotted outlines mark the adult γ lobes. Scale bar indicates 30*µ*m. **(J)** Ranking of pruning defects of hemispheres with and without transgene expression in the APL, within the experiments shown in A-I. Statistical significance calculated between hemispheres expressing or not in APL, using paired Wilcoxon signed-rank test with continuity correction, p=0.000133 for APL>Kir2.1, (C; n=31), p= 0.02473 for APL>TNT (F; n= 14), and p= 0.833 for APL>Kir2.1, 71QF2>Kir2.1 (I; n=12).

Next, we asked whether increasing APL activity is sufficient to induce early or more extensive pruning, we activated APL neurons using dTrpA1. Expressing dTrpA1 in APL neurons and activating them by rearing the flies in 29°C from 0 APF up to 18 APF, did not result in any change in the rate or extent of pruning of γ-KCs, as measured at the peak of remodeling (Figure S2 D-F, quantified in G). Taken together, these results suggest APL activity is required, but not sufficient, to hyperpolarize γ-KCs and to promote efficient axon pruning.

The major inhibitory neurotransmitter in the nervous system is GABA. GABA exerts its inhibitory function by binding to two types of receptors, the ionotropic GABA-A receptor, which conducts Cl^-^ upon ligand binding, and the metabotropic (G protein-coupled) GABA-B receptor, which induces K^+^ ion conductance as well as reduces Ca^2+^ current through voltage dependent Ca^2+^ channels (Gahwiler and Brown, 1985; Janigro and Schwartzkroin, 1988; Menon-Johansson et al., 1993). While the ionotropic GABA-A receptors have been shown to be excitatory during early pupal development and only become inhibitory during late development (Ryglewski et al., 2017), the metabotropic GABA-B receptors are thought to be hyperpolarizing throughout life (Cherubini et al., 1991). The *Drosophila* GABA-A receptor, *rdl*, is expressed in γ-KCs, however its expression level sharply decreases just prior to pruning in WT animals (Figure S3 A, Alyagor et al., 2018). We followed the expression of GABA-B-R1 using a protein-GFP fusion (GABA-B-R1^GFP)^ generated by means of Minos Mediated Integration Cassette (MiMIC; Bellen et al., 2015; Venken et al., 2011a). Interestingly, we detected GABA-B-R1-GFP in the MB calyx, the dendritic region of the KCs, before, during, and after metamorphosis (Figure S3 B-D). In order to verify that the GABA-B-R1 mimic line is specifically expressed in the KC calyx and to determine the cellular source of GABA-B-R1, we knocked down GABA-B-R1 expression by expressing RNAi using the strong pan-KC driver OK107-Gal4. Indeed, GABA-B-R1^GFP^ expression in the clayx was dramatically reduced, thus confirming the specificity of the RNAi, the GABA-B-R1^GFP^ reporter, and the fact that γ-KCs are the primary source of GABA-B-R1 in this brain region at the larval stage (Figure S3 E-F). Taken together, these results suggest that GABA signaling, probably via GABA-B-R1, could mediate hyperpolarization of γ-KCs at the initial stages of pruning. Indeed, knocking down GABA-B-R1 in all KCs using RNAi (driven by OK107-Gal4), or knocking it out in γ-KCs using tsCRISPR (by R71G10-Gal4-driven Cas9), both resulted in mild pruning defects (Figure S3 H-J quantified in G, using Mann Whitney U test, p-value =0.02444 for Irk1 RNAi n= 11 vs control n= 20 and < 0.00001 for tsCRISPR n=40 vs control n= 39).

**Figure S2.**
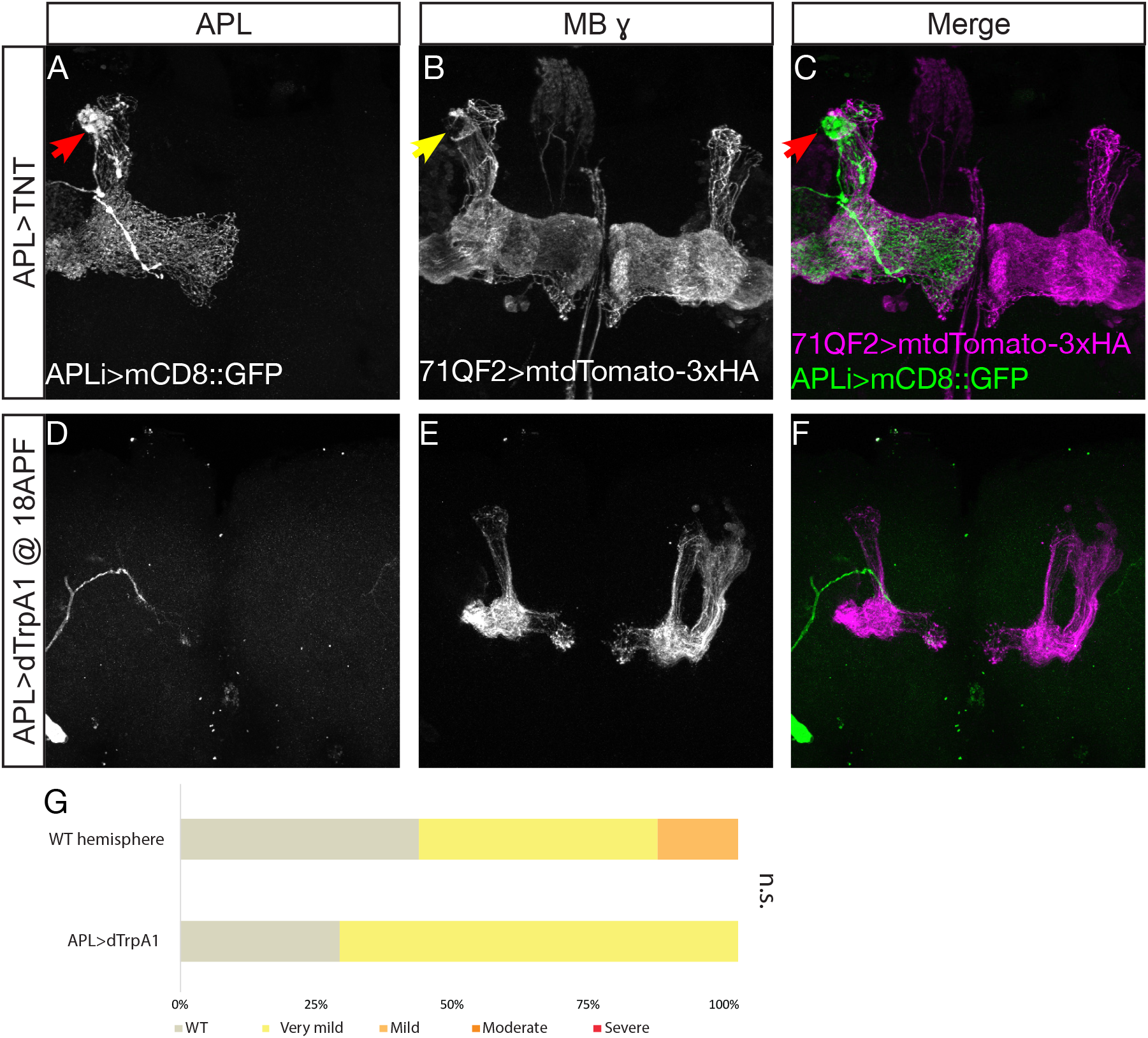
Expression of TNT in APL induces blebbing and activation of the APL is not sufficient to induce KC pruning. **(A-C)** Confocal Z projections of adult brains expressing mCD8::GFP and TNT under the control of APLi-Gal4 (grey in A and green in C) and mtdTomato-3cHA under the control of GMR71G10-QF2 (grey in B and magenta in C). Red arrowheads point to blebbing in the APL, yellow arrowheads point to unpruned γ-KC axons. **(D-F)** Confocal Z projections of brains of 18h APF pupa reared in 29^0^C from 0hAPF expressing mCD8::GFP and dTrpA1 under the control APLi-Gal4 (grey in D and green in F) and mtdTomato-3xHA under the control of GMR71G10-QF2 (grey in E and magenta in F). **(G)** Ranking of pruning defect severity in WT hemispheres (shown on the right in E and F) vs hemispheres expressing the dTrpA1 under the control of the APLi driver (D and F left hemispheres) grown in 29^0^C.

**Figure S3.**
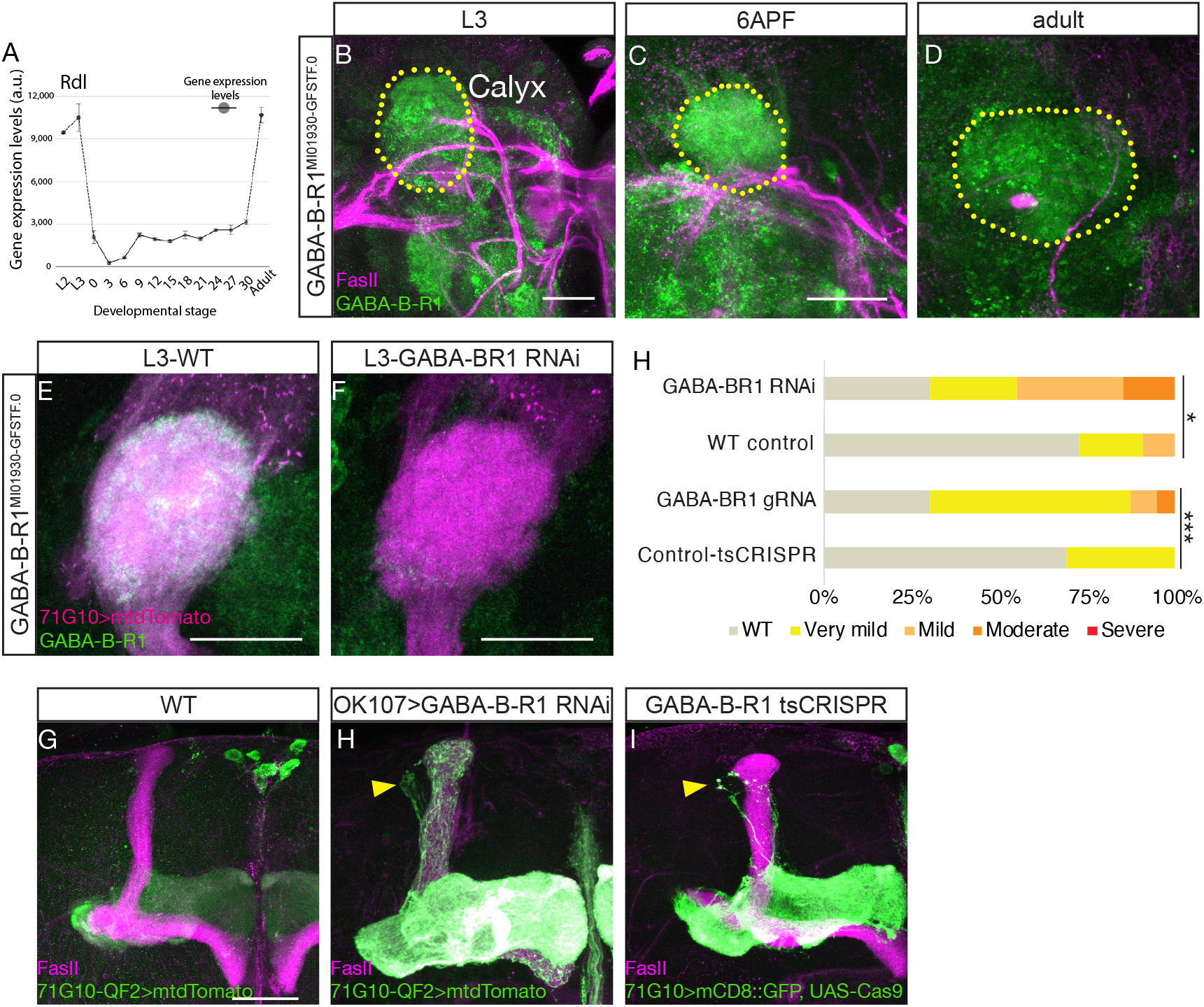
GABA BR1 expression is required for KC pruning. **(A)** Expression profile of Rdl in γ-KCs throughout development, showing an acute decrease in expression levels at the onset of metamorphosis. Extracted from (Alyagor et al., 2018), L2-L3 are 2^nd^ and 3^rd^ larval stages, 0-30 are hours APF.. **(B-D)** Confocal Z projections of L3 (B, E, F), 6h APF (C) and adult (D) brains expressing GFP-GABA-B-R1^MI01930-GFSTF.0^ (green; MiMiC lines from Bellen et al., 2015; Venken et al., 2011), immunostained with anti fasII (magenta). **(E-F)** GABA-B-R1^MI01930-GFSTF.0^ L3 brains expressing (E) mtdTomato-3xHA under the control of GMR71G10-QF2 (magenta) or additionally (F) GABA-B-R1 RNAi ^HMC03388^. **(G-I)** Confocal Z projections of adult brains immunostained with anti-FasII (magenta) expressing mtdTomato-3xHA under the control of GMR71G10-QF2 (green), brains additionally expressing GABA-B-R1 RNAi ^HMC03388^ (H) or UAS-Cas9.C and GABA-B-R1 gRNA (I) under the control of OK107-Gal4. **(J)** Ranking of pruning defects in (G-J). Statistical significance calculated using Mann-Whitney test, p-value =.02444 for GABA-BR1 RNAi (I; n= 11) vs control (H; n= 20) and < .00001 for tsCRISPR (J; n=40) vs control (not shown; n= 39). Scale bar indicates 30*µ*m.

### Irk1 and GABA-B-R1 are synergistically required for γ-KC pruning

The mild nature of the pruning defects induced by Irk1 knockdown or APL silencing, compared to the dTrpA1-induced pruning defect, could suggest that both processes work in parallel to hyperpolarize the γ-KCs prior to pruning. To investigate the involvement and possible epistasis of APL activity and irk1 we perturbed both genes in parallel. Interestingly, while the single gene perturbation of either irk1 or GABA-B-R1 resulted, as expected, in mild to moderate pruning defects (Figure 4 A-C, quantified in E, using Mann Whitney U test, p-value < .00001 for Irk1 RNAi n= 20 vs control n= 20 and 0.0001 for tsCRISPR n=20 vs control n= 20), the combined perturbation of both resulted in a significant phenotype exacerbation (Figure 4 D, Mann Whitney U test, p-value < 0.00001 for Irk1 RNAi combined with GABA-B-R1 tsCRISPR n= 20 vs control n= 20 or either irk1 RNAi or GABA-B-R1 tsCRISPR alone). These results suggest that the cell-autonomous effect of irk1 and APL neuronal activity work in concert to hyperpolarize γ-KCs at the onset of metamorphosis prior to pruning.

**Figure 4.**
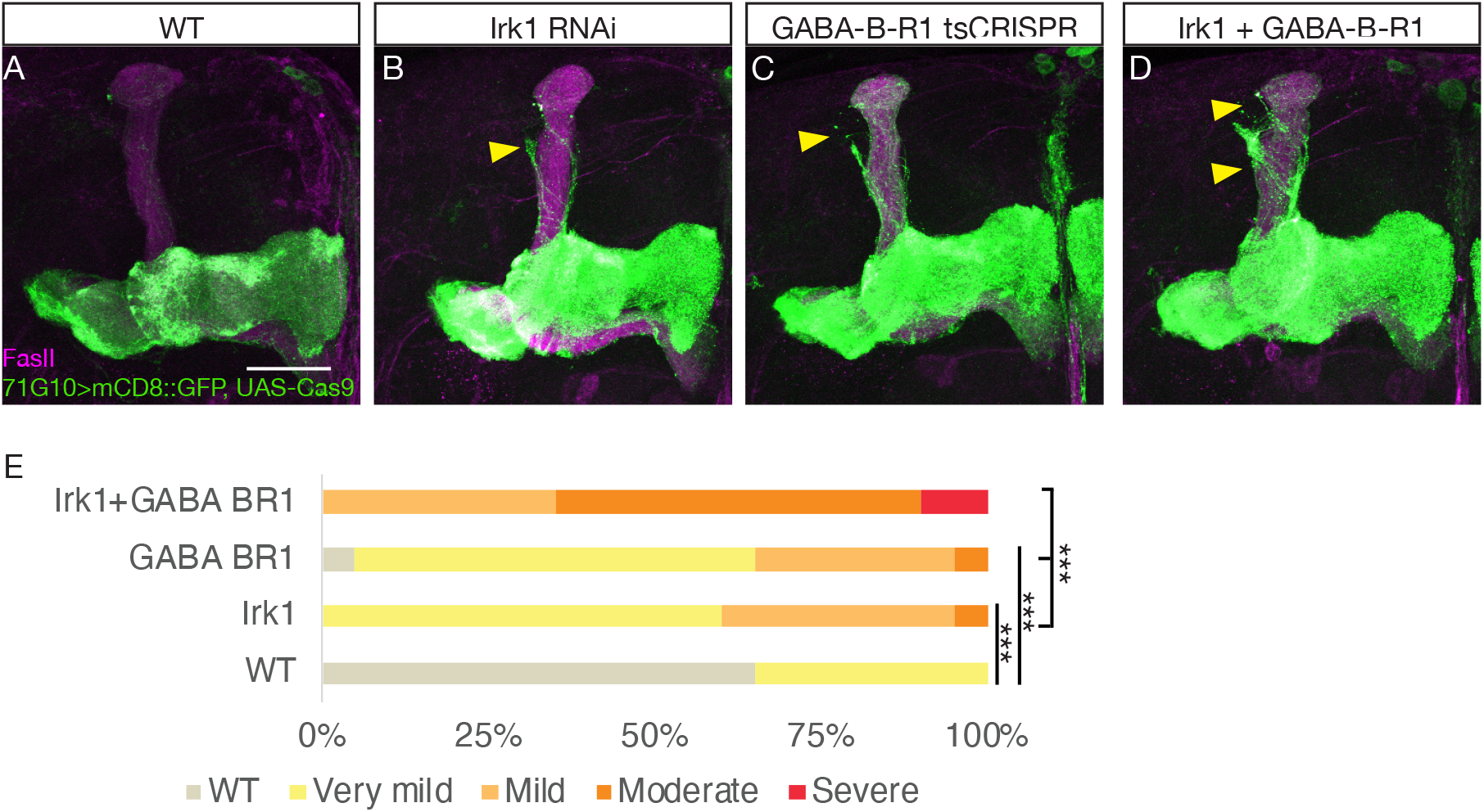
Irk1 and GABA-B-R1 expression are synergistically required for γ-KCs pruning. **(A-D)** Confocal Z projections of brains immunostained with anti-FasII (magenta), expressing mCD8::GFP and Cas9.C under the control of 2 copies of GMR71G10-Gal4 (A), additionally expressing UAS-Irk1-RNAi^HMS02480^ (B), U6.3:GABA-B-R1 gRNA (C), or both (D). Yellow arrowheads point to unpruned axons. Scale bar indicates 30*µ*m. **(E)** Ranking of pruning defect in A-D. Statistical significance calculated using Mann-Whitney test, p-value < .001 for Irk1 RNAi (B; n= 20) vs control (A; n= 20), p<0.001 for GABA-B-R1 tsCRISPR (C; n=20) vs control, and p < 0.001 for Irk1 RNAi combined with GABA-B-R1 tsCRISPR (D; n= 20) vs control or either vs irk1 RNAi or vs GABA-B-R1 tsCRISPR alone.

## Discussion

Precise neuronal circuitry is sculpted out of initial exuberant connectivity that is established during early development (Lee et al., 1999, 2000; Schuldiner and Yaron, 2014). This is achieved by the combination of both intrinsic and extrinsic factors (Meltzer and Schuldiner, 2020). Neuronal activity has been shown to play a major role in remodeling excitatory circuits, often via competition that is derived from experience-dependent or intrinsically generated activity waves (Hebb, 1949; Hensch, 2005; Katz and Shatz, 1996; Wong et al., 1993). However, the role and involvement of inhibitory synapses in activity-mediated plasticity is less clear. Likewise, the role of activity in stereotypic developmental remodeling, where competition is not apparent, is also not well understood. Here we demonstrate that silencing neuronal activity is a prerequisite for the initiation of pruning of the γ-KC axons. We show that this is achieved by the combination of cell-autonomous expression of irk1 along with inhibitory APL activity, likely signaling through GABA-B-R1 activation. Overall, we suggest that active KC post-synapses are stabilized, and that destabilization in the course of pruning requires silencing or hyperpolarization.

### Differential influence of Ca^2+^ dynamics and neuronal activity on pruning in the PNS vs CNS

Previously, Kanamori and colleagues have nicely demonstrated that transient compartmentalized Ca^2+^ influx through voltage gated Ca^2+^ channels activate Ca^2+^ dependent proteases to allow for the normal progression of pruning in class IV dendritic arborization (C4da) neurons (Kanamori et al., 2013). In contrast, activation of γ-KCs and subsequent entry of Ca^2+^ inhibits pruning. Similarly, while hyperpolarizing C4da neurons inhibited their pruning, it seems that hyperpolarization is actually required for γ-KC pruning. Taken together, these differences highlight the context dependent nature of Ca^2+^ signaling, and could be consolidated when we consider the neuronal systems themselves. While in C4da neurons only the dendrites undergo remodeling, in γ-KCs both the dendrites and axons remodel. Moreover, γ-KCs lie in the heart of the CNS of the fly, while C4da neurons reside in the PNS. As such, the characteristics of their connectivity is drastically different in nature. While γ-KCs have many elaborate pre- and post-synaptic partners with the potential to influence KC excitation and subsequent cytoplasmic Ca^2+^ levels, C4da dendrites are somatosensory, and thus connect to the epithelium rather than other synaptic partners. Moreover, it has been recently demonstrated that the ENaC (endothelial sodium channel) Pickpocket 26 (Ppk26), is actively degraded in C4da neurons (Krämer et al., 2019), suggesting that C4da actively reduce their excitability prior to pruning. Thus, the precise roles of excitability and Ca^2+^ levels during remodeling of axons and dendrites need to be further clarified.

### Neuronal excitability as a regulator of neuronal remodeling

The current concept of neuronal activity-mediated plasticity is focused on the plasticity of excitatory connections, and is generalized as ‘use it or lose it’. This is commonly interpreted such that connections with stimulated (or correlated) inputs grow stronger, while connections with inactive (or uncorrelated) inputs grow weaker. This process is based on mechanisms underlying Hebbian plasticity (Hebb, 1949). Hensch and colleagues have formulated a compelling hypothesis including inhibition in the formation of critical periods (Ferster, 2004; Hensch, 2005; Hensch and Fagiolini, 2005). In this model, lateral inhibition modulates Hebbian-type plasticity by enhancing the correlative activities of adjacent cortical neurons and producing anti-correlative activities in distal cells. Their model suggests that incorporation of GABAergic inhibition, downstream of retinal input, can provide a scaffold for the mature circuit. Interestingly, our findings that persistent γ-KC synapses inhibit APL pruning (Mayseless et al., 2018), and that APL activity is necessary for γ-KC pruning, are consistent with this model. As such, in a similar manner to the visual system of mammals, manipulation of GABAergic feedback inhibition modulates the excitatory and corelated state of the remodeling neurons, and influence their subsequent refinement.

Altogether, the *Drosophila* mushroom body, with its wealth of available genetic tools and well-characterized connectivity, can now be used to ask questions which were up to now inaccessible. For example, the relative contribution and influence of neuromodulatory neurons, such as dopaminergic and serotonergic neurons, on γ-KC remodeling and vice versa, and how this coordination subsequently influences the mature circuit and behavior, can now be assayed. Therefore, the mushroom body opens a route to extend our knowledge about the mechanistic epistasis of how mature circuit connectivity arises.

## KEY RESOURCE TABLE

| REAGENT or RESOURCE | SOURCE | IDENTIFIER |
| --- | --- | --- |
| **Antibodies** |  |  |
| Chicken anti GFP 1:500 | AVES | GFP-1020<br>AB 10000240 |
| Mouse monoclonal anti FasII 1:25 | Developmental Studies Hybridoma Bank (DSHB) | 1D4<br>AB 528235 |
| Rat monoclonal anti HA 1:250 | Sigma Aldrich | 11867423001<br>AB 10094468 |
| FITC Goat anti chicken 1:300 | Invitrogen | A-16055<br>AB 2534728 |
| Alexa fluor 647 Goat anti rat 1:300 | Invitrogen | A-21247<br>AB 141778 |
| Alexa fluor 647 goat anti mouse 1:300 | Invitrogen | A-32728<br>AB 2633277 |
| **Experimental Models: Organisms/Strains** |  |  |
| *D. melanogaster*: w[*]; P{y[+t7.7] w[+mC]=10XUAS-IVS-mCD8:GFP}attP2 | Gift from (Aso et al., 2014) | N/A |
| *D. melanogaster*: w[*]; P{y[+t7.7] w[+mC]=10XUAS-IVS-mCD8:GFP}attP40 | Bloomington drosophila stock center (BDSC) | BDSC: 32185;<br>FlyBase:<br>FBst0032185 |
| *D. melanogaster*: NP2631-Gal4, GH146-flp/CyO ; Mkrs,Sb/Tm6,Hu,Tb | Bloomington drosophila stock center (BDSC) | BDSC: 32186;<br>FlyBase:<br>FBst0032186 |
| *D. melanogaster*: TubP,FRT,Gal80,FRT,UAS-mCD8:GFP/CyO ; Mkrs,Sb/Tm6,Hu,Tb | Gift from (Lin et al., 2014) | N/A |
| *D. melanogaster*: GMR71G10-QF2<sup>Hack</sup> | Bloomington drosophila stock center (BDSC) | BDSC: 6872;<br>FlyBase:<br>FBst0006872 |
| *D. melanogaster*: UAS-TNT |  |  |
| *D. melanogaster*: OK107-Gal4 | Bloomington drosophila stock center (BDSC) | BDSC: 854; FlyBase:<br>FBti0004170 |
| *D. melanogaster*: QUAS-mtdTomato-3xHA |  |  |
| *D. melanogaster*: GMR71G10-Gal4 | Bloomington drosophila stock center (BDSC) | BDSC: 39604;<br>FlyBase:<br>FBti0137964 |
| *D. melanogaster*: UAS-TNT (w[*]; P{w[+mC]=UAS-TeTxLC.tnt}R3) | Bloomington drosophila stock center (BDSC) | BDSC: 28997;<br>FlyBase:<br>FBti0038570 |
| *D. melanogaster*: UAS-Kir2.1 (w[*]; P{w[+mC]=UAS-Hsap\KCNJ2.EGFP}7) | Bloomington drosophila stock center (BDSC) | BDSC: 6595;<br>FlyBase:<br>FBti0017552 |
| *D. melanogaster*: QUAS-Kir2.1 | Bloomington drosophila stock center (BDSC) | BDSC: 91802 |
| *D. melanogaster*: UAS-dTrpA1 | Bloomington drosophila stock center (BDSC) | BDSC: 26264;<br>FlyBase: FBti0114502 |
| *D. melanogaster*: UAS-Irk1-RNAi<sup>HMS02480</sup> | Bloomington drosophila stock center (BDSC) | BDSC: 42644;<br>FlyBase: FBti0151128 |
| *D. melanogaster*: UAS-GABA-B-R1-RNAi<sup>HMC03388</sup> | Bloomington drosophila stock center (BDSC) | BDSC: 51817;<br>FlyBase: FBti0157782 |
| *D. melanogaster*: w[1118]; P{y[+t7.7] w[+mC]=UAS-Cas9.C}attP2 | Bloomington drosophila stock center (BDSC) | BDSC: 54595;<br>FlyBase: FBti0159186 |
| *D. melanogaster*: GABA-B-R1 gRNA (y[1]; M{v[+t1.8]=WKO.1-G2}ZH-86Fb) | Bloomington drosophila stock center (BDSC) | BDSC: 83036;<br>FlyBase: FBti0205338 |
| *D. melanogaster*: Irk1 gRNA (y[1] v[1]; P{y[+t7.7] v[+t1.8]=U6.2-Irk1.dgRNA}attP40) | Bloomington drosophila stock center (BDSC) | BDSC: 92531;<br>FlyBase: FBti0215651 |
| *D. melanogaster*: w[*]; P{y[+t7.7] w[+mC]=UAS-CaMPARI.V398D}attP40 | Bloomington drosophila stock center (BDSC) | BDSC: 58762;<br>FlyBase: FBti0166951 |
| *D. melanogaster*: w[1118] P{y[+t7.7] w[+mC]=UAS-CaMPARI2.L398T} su(Hw)attP8 | Bloomington drosophila stock center (BDSC) | BDSC: 78320;<br>FlyBase: FBti0199492 |
| *D. melanogaster*: y[1] w[67c23]; Mi{PT-GFSTF.0}GABA-B-R1[MI01930-GFSTF.0] | Bloomington drosophila stock center (BDSC) | BDSC: 60522;<br>FlyBase: FBti0178464 |
| **Software and Algorithms** |  |  |
| FIJI | Image J | https://imagej.net/Fiji/Downloads |
| **Other** |  |  |
| Zeiss LSM 710 and 800 confocal microscope |  |  |
| 40x 1.3 NA oil immersion lens | Zeiss |  |
| TCS SP8 confocal microscope | Leica |  |

## CONTACT FOR REAGENT AND RESOURCE SHARING

Further information and requests for resources and reagents should be directed to and will be fulfilled by the Lead Contact, Oren Schuldiner

## EXPERIMENTAL MODEL

### *Drosophila melanogaster* rearing and strains

All fly strains were reared under standard laboratory conditions at 25°C (unless stated otherwise) on molasses containing food. Males and females were chosen at random. Developmental stage is referred to in the relevant places while adult refers to 3-5 days post eclosion.

APLi lines: NP2631-Gal4, GH146-flp/CyO; Sb/Tm3,Ser and TubP,FRT,Gal80,FRT,UAS-mCD8:GFP/CyO; Sb/Tm3,Ser, were generated and kindly provided by Dr. Gero Miesenböck (Lin et al., 2014).

71G10-QF^Hack^ was kindly provided by Dr. Christopher J. Potter (Lin and Potter, 2016).

## METHOD DETAILS

### Immunostaining

*Drosophila* brains were dissected in cold ringer solution, fixed using 4% paraformaldehyde (PFA) for 20 minutes at room temperature (25^0^C) on a nutator, after which brains were washed several times in PBT (phosphate buffer supplemented with 0.3% triton-x) blocked using heat inactivated goat serum and subjected to primary antibody staining overnight at 4^0^C, followed by three washes with PBT, then staining with secondary antibodies for 2 hours at RT, secondary antibodies were quickly washed with PBT and then washed again 3 times.

### Imaging and image processing

All stained brains were mounted on Slowfade (Invitrogen) and imaged on Zeiss LSM 710 or 800 confocal microscopes using 40x 1.3 NA oil immersion lens. Images were processed with ImageJ (NIH).

### Ranking

Ranking was performed on maximum Z-projections by one or more independent scorers in a double-blind manner with similar significance scores. The results of one scorer are shown for simplicity. Examples of rank severity score are shown in Figure S1 (A-E).

### Larval staging

Late vs early 3^rd^ instar (L3) larval stage was set by rearing larvae on fly food with added red food coloring. Early L3 larvae displayed a colored gut, while late L3 larvae had no coloration.

### CaMPARI photoconversion and imaging

Drosophila of the appropriate genotype were collected at the marked developmental stage; Early L3 (wandering stage with coloration in gut), Late L3 (wandering stage with no gut coloration), 0h APF (white pupa), 3h APF, and 6h APF. Larvae/pupae were illuminated using a UV illumination table (395nm) for 15 minutes, directly dissected in ice cold Ca^2+^ free Ringers solution, and immediately mounted on an aluminum foil wrapped microscope slide for imaging using a 20x Objective with 2.5 Zoom on an SP8 Leica confocal microscope. Hybrid detector 1, between 510nm and 545nm; and Hybrid detector 2, between 568nm and far red (maximum), averaging 2 iterations per line. Two different wavelength lasers were used: 488nm (10% intensity) and 561nm (10% intensity) scanned simultaneously. It was confirmed beforehand that there was no “spill-over” between detection channels.

ImageJ was used to measure the fluorescence in the acquired images in selected ROIs, which had background subtracted using an identical ROI measuring background intensity. Red/Green fluorescence was normalised to the mean of the first time-window (divided every measurement by the mean of the first group).

### Quantification and statistical analysis

In all cases, *** represent a p-value lower than 0.001; ** represent a p-value lower than 0.01 and * represents a p-value lower than 0.05. Statistical tests were run using R-Studio or OriginPro 8.5G. specific p-values and sample sizes and testes are indicated in the relevant figure legend and in text.

## Acknowledgements

We thank C. Potter, GM. Rubin, the Kyoto (DGRC) and Bloomington Stock Centers for reagents; H. Meltzer, for graphical illustrations, discussions, critical reading, support and other members of the Schuldiner lab, especially I. Alyagor, N. Kollet, S. Yaniv, and V. Berkun, for discussions, critical readings of the manuscript and overall support and assistance during the different working stages. This work was supported by Volkswagen Stiftung (joint Lower Saxony – Israel) grant #A112379 to A.F. and O.S., the Israel Science Foundation (ISF) grant #1100/16 to O.S., the German Research Foundation (FOR 2705) to A.F., and an EMBO short term fellowship #382-2016 to O.M.

## Author contributions

O.M. designed, performed, and analyzed experiments, wrote the manuscript and procured funding; R.E.Y. and G.S. performed and analyzed experiments; A.F. designed CaMPARI experiments, assisted in data interpretation and analysis, critically read and discussed the manuscript, and procured funding; O.S. designed and analyzed experiments, wrote the manuscript, and procured funding.

## Declaration of Interests

The authors declare no competing interests.

## Notes

### Competing Interest Statement

The authors have declared no competing interest.

